# Memory loss at sleep onset

**DOI:** 10.1101/2022.04.25.489361

**Authors:** Célia Lacaux, Thomas Andrillon, Isabelle Arnulf, Delphine Oudiette

## Abstract

Every night, we pass through a transitory zone at the borderland between wakefulness and sleep, named the first stage of non-rapid eye movement sleep (N1). N1 sleep is associated with an increased hippocampal activity and dreamlike experiences that incorporate recent wake materials, suggesting that it may be associated with memory processing. Here, we investigated the specific contribution of N1 sleep in the processing of memory traces. Participants were asked to learn the precise locations of 48 objects on a grid and were then tested on their memory for these items before and after a 30-minute rest during which participants either stayed fully awake, transitioned toward N1 or deeper (N2) sleep. We showed that memory recall was lower (10% forgetting) after a resting period including only N1 sleep compared to N2 sleep. Furthermore, the ratio of alpha/theta power (an EEG marker of the transition towards sleep) correlated negatively with the forgetting rate when taking into account all sleepers (N1 and N2 groups combined), suggesting a physiological index for memory loss that transcends sleep stages. Our findings suggest that interrupting sleep onset at N1 may alter sleep-dependent memory consolidation and promote forgetting.

## INTRODUCTION

Numerous human studies have demonstrated that sleep improves or stabilizes memory in a variety of tasks, including perceptual, associative, spatial, motor, and emotional learning (Born et al., 2006; Diekelmann & Born, 2010; Paller et al., 2021; Rasch & Born, 2013). A growing body of research on animals supports the notion that this sleep-dependent memory consolidation is enabled by a hippocampo-neocortical dialogue orchestrated by slow oscillations during NREM sleep (Maingret et al., 2016; Ólafsdóttir et al., 2018). Importantly, sleep does not blindly consolidate all memories formed during the day, but rather selectively consolidates those that are expected to be of future relevance (Oudiette et al., 2013; Saletin et al., 2011; Stickgold & Walker, 2013). How the sleeping brain performs this ‘memory triage’ remains mysterious (Stickgold & Walker, 2013). One could imagine that the first sleep stage, known as NREM sleep stage 1 (N1 sleep), could contribute to this process by reviewing recent experiences and either deleting the memory traces deemed irrelevant (if the tagging process occurred pre-sleep), or directly tagging the memory traces considered important for consolidation in subsequent sleep stages. However, because animal models used in sleep research lack an equivalent to the classically described NREM subdivision in humans (Lacroix et al., 2018), they cannot provide information about the role of each NREM stage, particularly N1, in memory. On the other hand, cognitive researchers have paid little attention to the N1 stage, possibly due to its fleeting nature, leaving us in the dark about its potential role in memory processing.

However, several factors suggest the involvement of sleep onset in memory processing. First, the hippocampus, a key brain region for memory consolidation, exhibits increased activity in the late N1 period compared to wakefulness (Picchioni et al., 2008). Second, N1 is associated with vivid, dream-like experiences (named ‘hypnagogia’) that often integrate recent daytime events (Stickgold et al., 2000; Wamsley, Perry, et al., 2010; Wamsley, Tucker, et al., 2010; Wamsley & Stickgold, 2019). One could hypothesize that this reviewing of recent events during hypnagogic experiences reflects the initial steps of memory processing, such as the tagging of memories for consolidation in subsequent NREM sleep (Stickgold, 2009). In support of this idea, memory-related experiences that are incorporated into sleep onset mentation are later found in the contents of NREM and REM sleep dreams within the same night of sleep (Wamsley & Stickgold, 2019). Additionally, several studies have shown a positive correlation between dreaming about a task and subsequent memory performance (De Koninck et al., 1990; Plailly et al., 2019; Wamsley, Tucker, et al., 2010; Wamsley & Stickgold, 2019). Combined, such findings would imply that the same memories are processed sequentially across the successive sleep stages (reminiscent of the ‘sequential hypothesis’, Giuditta, 2014), with N1 serving as an initiating stage, tagging. This idea is substantiated by Stenstrom et al. (2012) who found that hypnagogic experiences connect fragments from distal memory sources that share semantic similarity, suggesting that a hippocampal-dependent integrative process (Shohamy & Wagner, 2008) occurs during this stage (i.e., the blending of overlapping past events into an integrated memory representation). Recently, we confirmed the role of N1 in such a gist abstraction process (Lacaux et al., 2021). We found that spending on average one minute in N1 tripled the chance of discovering a hidden regularity within memory traces (83%), compared to wakefulness (30%) or N2 sleep (14%).

However, beyond these findings, the literature on the sleep-onset period and memory remains sparse and inconclusive. One study showed that a 6-min nap was sufficient to improve word recall compared to a similar period of wakefulness (Lahl et al., 2008). However, those naps included both N1 and N2 sleep, thus preventing us from drawing a definitive conclusion about the respective contribution of each stage to the observed memory benefit. In contrast, older studies reported evidence that sleep onset was associated with retrograde amnesia of the materials encoded in the few minutes prior to sleep onset (Wyatt et al., 1994, 1997). In those studies, word pairs were presented to participants as they were falling asleep, so it is unclear whether sleep onset interfered with the encoding or consolidation of the stimuli.

Here, we aimed to better understand whether the twilight zone between wakefulness and sleep contributes to memory consolidation. To do so, we compared how a 30-minute resting period that included either only wakefulness, only N1, or N1+N2 sleep impacted the fate of memories that had been successfully encoded prior to the resting period.

## METHODS

### Participants

52 healthy volunteers (49% females, 24.83 ± 4.45 years) participated in the study. They were screened for exclusion criteria such as excessive daytime sleepiness, as well as history of sleep, neurological, or psychiatric disorders. To facilitate sleep onset, we asked participants to sleep about 30% less than usual during the night preceding the experiment and to avoid stimulants (e.g., coffee, tea) on the day of the experiment. Subjects were paid 10€ per hour as compensation for their participation (plus a bonus based on their performance). All subjects provided their written informed consent prior to the onset of the study. The study protocol was approved by the local ethics committee (Comité de Protection des Personnes, Ile-de-France III, Paris, France).

### Task

Subjects had to learn the precise location of 48 visual stimuli on a 8×6 squared grid background displayed on a 34.4-cm computer screen. Locations were randomly determined for each stimulus and each participant at the beginning of the session. Visual stimuli were pictures representing a variety of objects or animals and had a dimension of 156×156 pixels (corresponding to 4.13×4.13 cm). Subjects were instructed to reposition the pictures as precisely as possible in order to maximize their bonus monetary reward (maximum reward per picture = 10 cents). A black and white square indicated the center of the object and served as a reference point for the distance calculation. The *Psychloolbox* (Brainard, 1997; Kleiner et al., 2007) was used to program the task in Matlab R2018b. We computed two measures of accuracy for each object: 1) a continuous, precise measure consisting of the geometric distance between the position given by the participants and the correct position, and 2) a binary correct vs. incorrect measure. An object was considered correctly located if the participant placed it within 5.4 cm of its original location (corresponding to the diagonal of a square unit within the grid; see **Figure 1** for an illustration of the correct perimeter). This threshold was explicitly stated to participants and was the same as in Oudiette et al. (2013) and in Rudoy et al. (2009).

**Figure 1:**
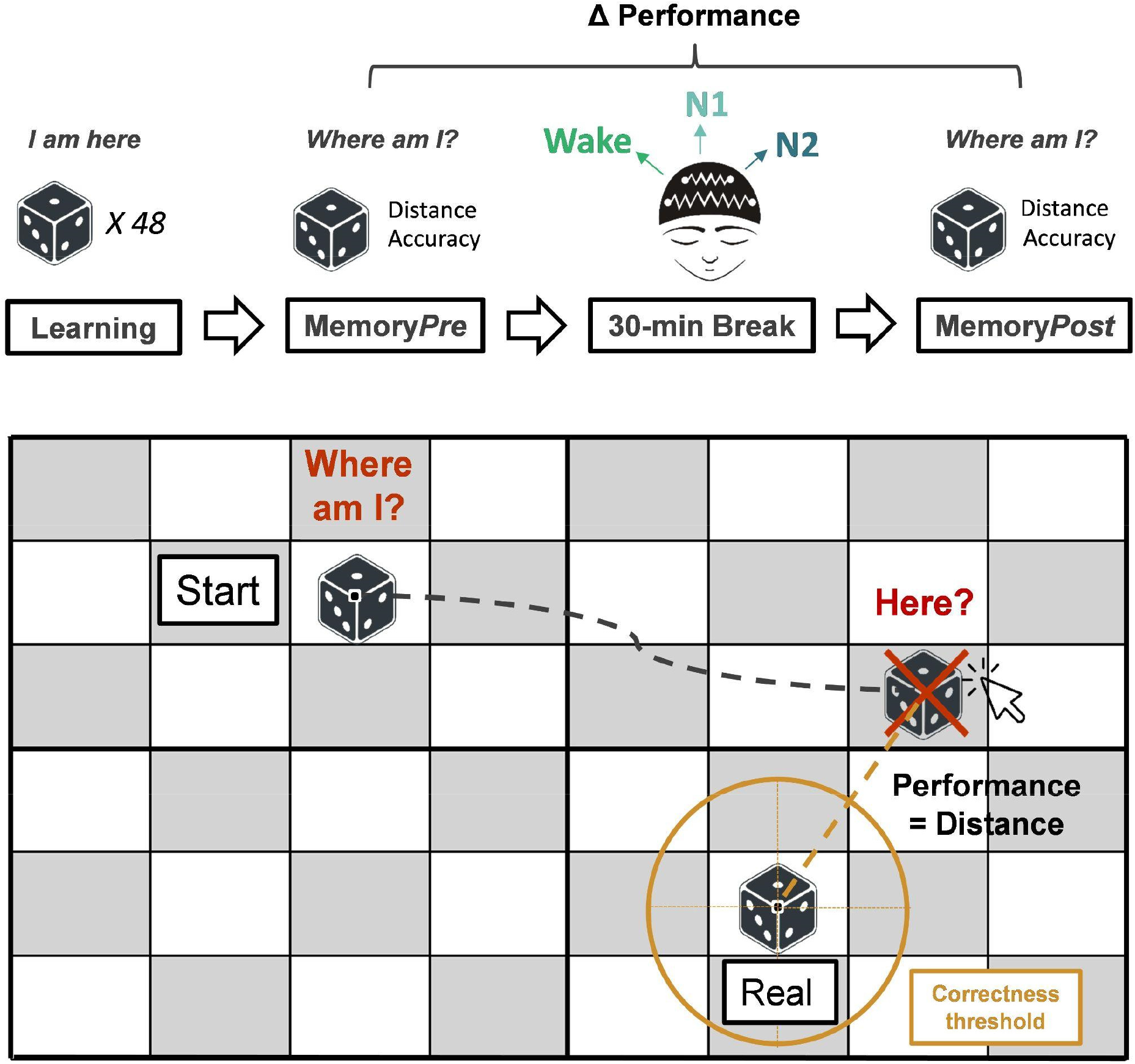
Experimental paradigm. (Top) Protocol timeline. Subjects first learnt the precise location of 48 pictures (e.g., a dice) which were presented within a grid. They were tested on their memory for these objects’ locations before and after a 30-min resting period, during which they could sleep while being continuously monitored via polysomnography. They were regularly awakened (approx. every 6 minutes) by a sound to prevent them from sleeping too deeply and to probe their mental content. Based on their sleep-wake states during the resting period, participants were divided into three groups: Wake (N = 16), N1 sleep (N = 15) and N2 sleep (N = 21). **(Bottom) Visuo-spatial memory test**. Each object appeared at a wrong position (start) and the subjects had to drag it to its correct (real) location as precisely as possible. Here, the object would be considered incorrect as the location provided by the subject is outside the circle ‘sperimeter, which represents the thresholdfor correctly-placed objects (see **Methods** for more details).

### Experimental design

Participants performed this spatial memory task (see **Figure 1** for details on the task) before and after a 30-min resting period while monitored with polysomnography (**Figure 1**). The protocol was divided into four main phases (summarized in **Figure 1**):

#### Learning phase

Participants first went through a **Learning phase** in which they had to memorize the precise location of 48 stimuli. First, the stimuli appeared one by one at the correct location, and each stimulus was repeated twice in a random order (passive viewing). The stimuli then appeared at a wrong location, and subjects had to drag them to the correct location using a computer mouse (active learning phase). They received feedback on their performance during this phase to ensure proper encoding. A picture was considered ‘encoded’ if the subject placed it correctly twice in a row (see the Task section for details on how accuracy is measured). The entire learning phase (passive viewing + active learning) was divided into three blocks of 16 stimuli, separated by a short break. It stopped when participants encoded the pictures’ positions well enough (i.e., 80% of correct stimuli). After completing the learning phase, participants were given a short, 3-min break before proceeding to the next phase *(Pre)*.

#### Phase 2 (Pre)

Participants were tested on their memory for the 48 pictures (**MemoryTest *Pre***). During the test, each object (one at a time) appeared in a wrong position, and participants had to drag the object to its correct location using a computer mouse, just like in the learning phase, but without feedback this time. Of note, unbeknownst to the participants, the starting position of an object was not determined randomly, but rather according to a hidden rule: it was always located on a fixed-length diagonal to the final/correct object’s location. This paradigm was originally designed to assess the impact of N1 on insight (i.e., the sudden discovery of a solution to a problem, here a hidden rule that allowed predicting the position of any object) too. But here, only participants who did not find the rule (N = 52) were considered, and the part on insight will be the subject of a separate report.

#### Phase 3 (Break)

After the first memory test, participants were given a 30-minute resting period. They lay on a bed in a dark room, eyes closed, with the instruction to rest and sleep if they wished. Every 6 minutes (plus a jitter randomly chosen from 0 to 30 seconds), we played an awakening sound to wake participants up. After each awakening, participants were instructed to speak aloud what was going through their minds (e.g., thoughts, images, reveries, dreams) before the alarm. This procedure was aimed first at minimizing the probability that participants would fall into deeper sleep stages and thus maximizing the probability of obtaining resting periods with N1 as the only sleep stage. Second, it sought to assess the impact of spontaneous mental content on memory processing. According to this design, we later split participants into three distinct groups based on their sleep-wake state during the break (see demographic and sleep parameters in **Table 1**): a ‘Wake group’ (subjects who stayed awake during the whole break; N = 16), a N1 group (subjects who reached N1 without any signs of deeper sleep stages; N = 15), and a N2 group (subjects who reached N2 stage; N = 21, of which two directly transitioned from wakefulness to N2 and therefore without any N1). Critically, all experimental conditions were identical between these three groups.

**Table 1.** Demographic and sleep characteristics of each group. Measures are presented as the mean ± the standard deviation or in percentages for proportions. N_Wake_ = 16, N_N1_ = 15, N_N2_ = 21. Educational level was scored according to the International Standard Classification of Education (Schneider, 2013). P-values are shown (ANOVA or Kruskal-Wallis Tests when appropriate: ordinal data or violations of normality assessed with the Shapiro-Wilk test, and Chi-squared Test for comparisons of proportions). When appropriate, post-hoc comparisons have been computed with Tukey-Kramer correction for multiple comparisons, and we report significant differences between Wake and N1/N2 (*), N2 and Wake/N1 (#), and between all groups ($).

| Group | Wake | N1 | N2 | P<sub>value</sub> |
| --- | --- | --- | --- | --- |
| **Demographic data** |  |  |  |  |
| Age, years | $26.50 \pm 4.50$ | $24.10 \pm 3.83$ | $24.10 \pm 4.68$ | 0.20 |
| Laterality (right-handed), % | 100 (16/16) | 93.33 (14/15) | 85.71 (18/21) | 0.27 |
| Gender, women % | 62.50 (10/16) | 40 (6/15) | 71.43 (15/21) | 0.16 |
| Educational level (0-7) | $6.75 \pm 0.45$ | $6.67 \pm 0.49$ | $6.71 \pm 0.46$ | 0.88 |
| Epworth score (0-24) | $7.50 \pm 2.76$ | $8.0 \pm 3.02$ | $7.90 \pm 2.86$ | 0.87 |
| **Cofactors** |  |  |  |  |
| Distance Pre, cm | $1.38 \pm 0.38$ | $1.48 \pm 0.45$ | $1.20 \pm 0.31$ | 0.17 |
| Correct Pre, % | $82.55 \pm 10.72$ | $83.47 \pm 12.66$ | $86.90 \pm 8.96$ | 0.48 |
| PVT RT Pre, ms | $292.79 \pm 57.30$ | $270.89 \pm 18.98$ | $275.21 \pm 21.44$ | 0.72 |
| PVT RT Post, ms | $274.66 \pm 35.50$ | $268.46 \pm 24.49$ | $280.36 \pm 22.61$ | 0.36 |
| **Sleep measures** |  |  |  |  |
| Wake duration, min | $27.44 \pm 1.05$ | $23.26 \pm 3.82$ | $16.78 \pm 5.40$ | **<0.001<sup>\$</sup>** |
| N1 duration, min | $0 \pm 0$ | $4.40 \pm 3.75$ | $3.78 \pm 3.57$ | **<0.001<sup>\*</sup>** |
| N2 duration, min | $0 \pm 0$ | $0 \pm 0$ | $7.22 \pm 5.02$ | **<0.001<sup>#</sup>** |
| Sleep latency, min | NA | $13.53 \pm 7.06$ | $9.43 \pm 4.53$ | **<0.001** |

#### Phase 4 (Post)

After the resting period, participants were tested again on the memory task (**MemoryTest *Post***), using the same procedure as for the *Pre* test.

To measure potential differences in vigilance levels between groups, participants also performed a 3-min Psychomotor Vigilance Test (PVT) at the beginning of Phases 2 *(Pre)* and 4 (*Post*). The PVT consists of monitoring the appearance of a stimulus on a screen and responding to each appearance as fast as possible (Grant et al., 2017).

### Mental content

Each mental content reported during the resting period was labeled by the experimenters as a dream or thought, with each being further classified as task-related or not. Given that we were studying the wake-to-sleep transition, we opted for a more conservative definition of dream than is commonly used (i.e., any mental content during sleep). Here, a mental report was only deemed as a dream-like experience if it was ‘involuntary, spontaneous, perceptual, and bizarre’.

### Sleep monitoring

Subjects were monitored with video-polysomnography for the entire duration of the experiment. The montage included 3 EEG channels (FP1, C3 and O1), electro-oculograms (EOG) with electrodes placed on the superior and inferior outer canthi of the eyes, chin electro-myogram (EMG), a microphone, and infrared video recordings. The impedances of electrodes were generally below 5kΩ. EEG signals were referenced to A2 (right mastoid) and sampled at 250 Hz.

### Sleep scoring

EEG data was bandpass-filtered between 0.1 and 40 Hz and EOG derivations between 0.3 and 15 Hz (two-pass Butterworth filter, 5th order). EMG signal was obtained with a local derivation placed on the chin, which was bandpass-filtered between 10 and 100 Hz (two-pass Butterworth filter, 5th order). Participants’ sleep-wake states during the resting period were scored offline by two experienced scorers according to the standard sleep scoring guidelines of the American Academy of Sleep Medicine, AASM (Berry et al., 2012). There was a high concordance between these two independent scorers (CL and DO, Kappa coefficient > 0.8) and the remaining disagreements were examined by a third expert scorer (SL).

### EEG spectral analyses

A spectral decomposition of the pre-processed EEG signal was conducted on the entire resting period for the occipital O1 electrode. Epochs with an absolute amplitude above 500 μV were discarded from this analysis. Welch’s method was used to estimate the log-transformed power spectral density (PSD) for two frequency bands (Alpha: 8-12Hz and Theta: 4-7Hz) using a 6-s sliding-window with 50% overlap and a frequency resolution of 0.2Hz. The power over each 6-s window was averaged for each 30-s epoch. Epochs with an absolute amplitude greater than 150μV were excluded from this analysis. Finally, the corresponding power spectra were averaged across the whole break duration. A constant of 3 was added to the power values to guarantee that they were all greater than zero, before calculating an alpha/theta ratio for each subject. Of note, the reported findings are similar for constant values superior or equal to 3. We also computed a normalized version of the alpha/theta ratio by z-scoring the individual values within each group (Wake, N1, and N2 sleep).

### Statistical analyses

As our aim was to investigate whether sleep-dependent memory consolidation starts as early as N1, we restricted our behavioral analyses to items that were sufficiently encoded before sleep. Accordingly, we excluded data from pictures incorrectly recalled (i.e., pictures that were positioned above the distance threshold of 5.4cm from the original location during the *Pre* test; mean = 7.40 ± 5.11 items excluded). Fisher Tests were used to test relationships between categorical variables. Kruskal-Wallis Tests (or One-Way ANOVAs for normally-distributed data) were conducted to test the impact of the group (Wake, N1, or N2) on performance. When appropriate, additional *post-hoc* comparisons with Tukey-Kramer correction for multiple comparisons were performed. Wilcoxon signed-rank Tests were used to compare two paired non-parametric variables, and Mann-Whitney for independent samples. In the case of non-significant results, Bayesian statistics were computed using JASP (JASP Team, 2019) with a prior distribution following a Cauchy distribution with a default scale rate of 0.707. A Bayes Factor BF01 typically above 3 (i.e., the null hypothesis is three times more likely than the alternative hypothesis) provides supportive evidence for the null hypothesis.

Correlations between alpha/theta ratio and memory performance were performed using Pearson’s correlation (and correlation plotting using the *gramm* toolbox; Morel, 2018). The Cohen’s ? test was used to evaluate inter-judge agreement. All tests were two-tailed, and a probability level of less than 0.05 was considered significant. All computations were performed using Matlab, version 2018b (The MathWorks Inc®).

Finally, to estimate the respective influence of the subject groups (wake, N1, and N2) and alpha/theta ratios on the forgetting rate, we fitted a series of models with the forgetting rate as a predicted variable. The models were as follows:

Model 0: Forgetting-Rate ~ 1
Model 1: Forgetting-Rate ~ 1 + Group
Model 2: Forgetting-Rate ~ 1 + Group + Alpha/Theta
Model 3: Forgetting-Rate ~ 1 + Group * Alpha/Theta

These models were fitted using the *glm* function from the statistical package in R software. Model comparison was performed using the anova function and F-tests are reported in the **Results** section. *Post-hoc* comparisons were performed on the winning model and also using a F-test.

## RESULTS

Participants performed a memory task before and after a 30-min break during which they were allowed to sleep (see **Figure 1** and ***Methods*** for details on the experimental task and timeline). Throughout the break, subjects were regularly awakened (approximately every 6 minutes) to prevent them from falling too deeply into sleep and to probe their mental content. Depending on their sleep-wake state during the resting period, participants were later subdivided into three groups (Wake, N1, and N2; see the **Methods** for details).

Participants’ demographic and sleep parameters are provided in **Table 1**. The only difference between the Wake and N1 groups was the amount of time spent in N1 (respectively, 0 vs. 4.40 ± 3.75 min); subjects in the N2 group spent a similar amount of time in N1 than the N1 group (mean ± SD = 3.78 minutes ±3.57, z = 0.45, p = 0.65, Mann–Whitney U Test; BF_01_ = 2.86) plus an average of 7.22 (± 5.02) minutes in N2.

### N1 is associated with forgetting

We found that the change in recall accuracy (geometric distance) after the resting period significantly differed as a function of the group (Wake, N1, N2) (**Figure 2A**, χ^2^(2) = 6.82, p = 0.03, Kruskal-Wallis). The *Pre-Post* difference was lower than 0 (i.e., subjects worsened their performance compared to Pre) for all groups (p <0.001, Wilcoxon signed-rank tests), but this difference was larger for N1 subjects, who overall tended to place the objects further away from their correct location after the nap than before (mean Δ distance *pre-post:* Wake = −0.53 ± 0.42 cm, N1 = −0.83 ±0.89 cm, N2 = −0.27 ± 0.36 cm, p-value between N1 and N2 = 0.055; *post-hoc* comparisons with Tukey-Kramer correction for multiple comparisons). Results obtained with the binary (i.e., correct vs. incorrect) rather than continuous estimate of recall, were concordant, and N1 subjects tended to have a lower overall accuracy than Wake and N2 subjects (**Figure 2B**, mean Wake = −1.04%, N1 = −3.61%, N2 = + 0.10%, p = 0.088, Kruskal-Wallis Test).

**Figure 2.**
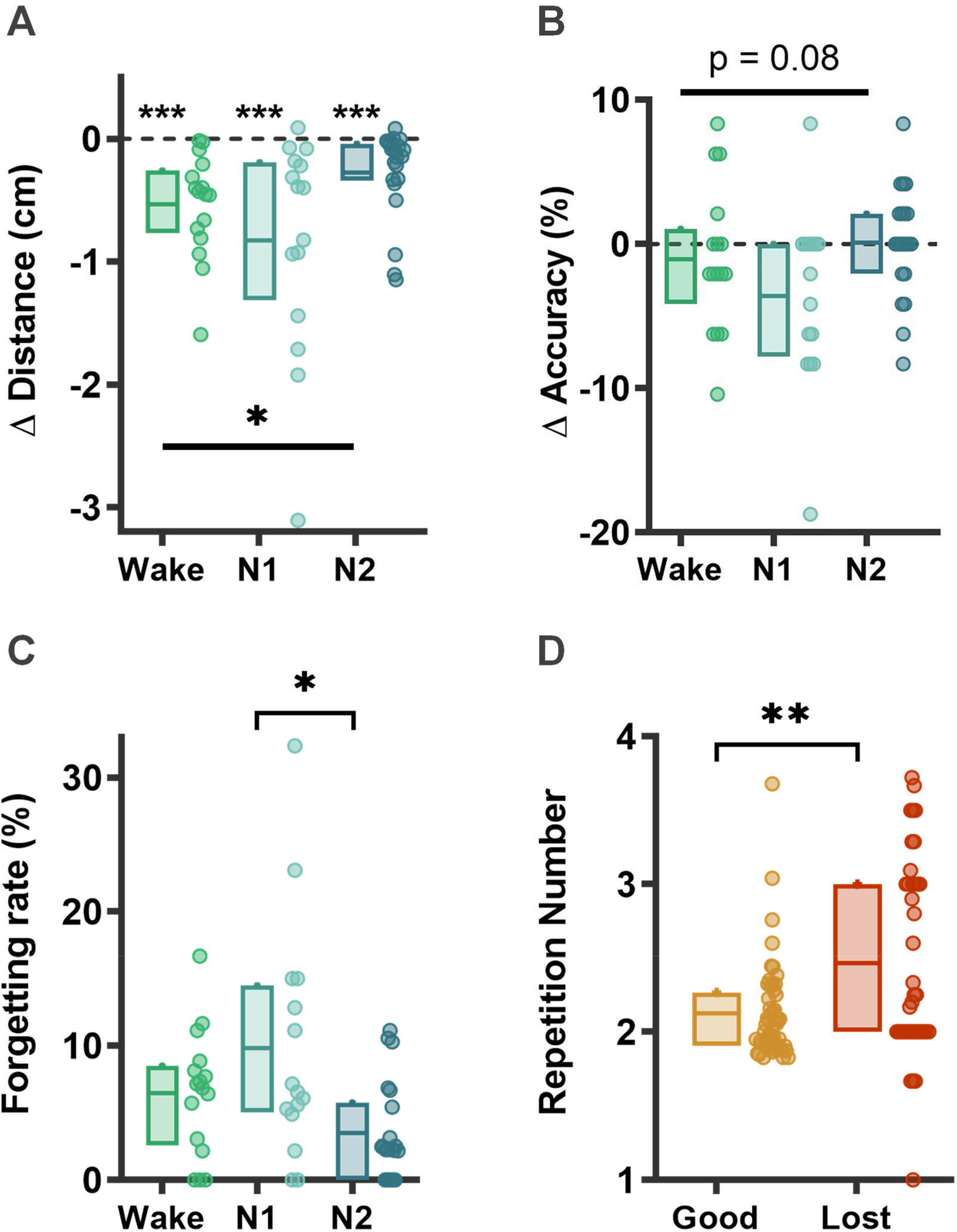
Memory loss at sleep onset. **(A)** Delta (pre-post) distance and **(B)** delta (post-pre) accuracy (percentage of correct objects) for each group (Wake, N1 and N2). Negative values thus indicate forgetting for both measures. The delta distance was computed on the items that were correct during the Pre test (items sufficiently encoded to potentially benefit from memory consolidation during the break). **(C)** Forgetting rate for each group: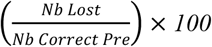. **(D)** Number of repetitions during the learning phase for the Always Good (correct in pre and post) and Lost items, all groups combined. For each box, horizontal lines represent the 1^st^, mean, and 3^rd^ quartiles. Individual data is also depicted by circles alongside their boxplot. Kruskal-Wallis tests were performed for comparisons between all three groups; when appropriate, post-hoc comparisons with Tukey correction have been computed. One-sample Wilcoxon signed-rank was used for comparison with zero level, and two-samples for comparisons between two paired samples: Always Remembered and Lost. Stars are used alone to report the p-values obtained when comparing data to zero, stars with a line for between-group differences, and stars above a square bracket for post-hoc comparisons. *** p<0.001; **, p<0.01; *, p<0.05,; ns, for nonsignificant differences.

Importantly, the overall decrease in performance observed in the N1 group was not due to baseline differences in memory abilities, as all groups performed equally well at the *Pre* phase both in terms of average distance from the correct location (p = 0.17, Kruskal-Wallis) and number of correctly placed objects (p = 0.48, see **Table 1**). Furthermore, control analyses indicate that the difference observed in the N1 group was not related to other confounding factors (e.g., sleep inertia), as all groups were not different in terms of sleepiness (Epworth sleepiness score), alertness (PVT score), or educational level (see **Table 1**). Of note, only 5 subjects out of 52 (9.62%) reported unambiguous task-related dreaming experiences, which does not allow statistical analyses on the role of mental content on memory performance. However, we collected a sufficient number of reported dreams in general or task-related thoughts to assess their impact on performance. There was no difference in the level of forgetting whether subjects reported dreaming experiences or not (mean forgetting rate in subjects with dream reports: 6.09% ± 4.54 vs without: 6.29% ± 8.61; p = 0.23, Wilcoxon rank sum test), nor if they thought about the task during the resting period (mean = 6.08 ± 5.94) or not (6.53 ± 6.90; p = 0.94, Wilcoxon rank sum test).

### Memory loss following N1 sleep concerns a subset of items

We then investigated whether this worsened performance was related to a general decrease in precision for all objects or if it was due to the forgetting of a subset of items. To do so, we classified the 48 objects into 4 categories: 1) Always Good (the remembered objects, correctly placed in both the *Pre* and *Post* phases), 2) Gained (falsely placed at *Pre*, correctly placed at *Post*), 3) Always Bad (the forgotten objects, falsely placed in both the *Pre* and *Post* phases), and 4) Lost (correctly placed at *Pre*, falsely placed at *Post*). Only the Lost objects category differed between groups (χ2(2) = 7.70; p = 0.02, Kruskal-Wallis Test; see **Supplementary Figure S1**). *Post-hoc* comparisons showed that there was a higher number of Lost objects (on average, 3.6 objects) following a resting period containing N1 sleep compared to one with N2 sleep (p-value between N1 and N2 = 0.02; p-value between Wake and N1 = 0.66; *post-hoc* comparisons with Tukey correction). This corresponded to a 10% forgetting rate 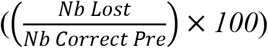, **Figure 2C**). Of note, this result also holds true when removing an outlier present in the N1 group (Kruskal-Wallis Test; p = 0.04).

Additionally, we attempted to determine what these lost items had in common. Wyatt et al. (1994) reported a sleep onset-related amnesia of stimuli presented just prior to sleep onset (3 minutes before), with no deficit for stimuli presented earlier. Here, we did not observe a difference in the timing of the presentation of these items, respective to sleep onset, between those that remained correct and those that were Lost during the active learning phase (3 possible blocks; mean presentation block Always-Good = 1.92, Lost = 2.06, p = 0.13) or during the *Pre* phase (48 stimuli; mean presentation order Always-good = 23.74, Lost = 25.49, p = 0.18, Wilcoxon-signed rank). Instead, we found that the Lost objects were the ones that participants had originally the most difficulty encoding (i.e., they were repeated more times during the Learning phase before encoding than the Always-good objects: **Figure 2D**; mean Lost = 2.46 repetitions, mean Always-good = 2.12; z = −2.74, p = 0.0061, Wilcoxon signed-rank Test; no between-group difference in the number of repetitions before encoding of these Lost and Always-good objects: p = 0.41 and p = 0.75 respectively).

### Neurophysiological substrate of memory loss

So far, we have categorized subjects according to standard sleep scoring methods. However, because it is based on discrete 30-second epochs, such classification misses subtle variations in electrophysiological activity (Hertig-Godeschalk et al., 2020; Hori et al., 1994) occurring at shorter timescales. To better understand the critical factors that may be associated with memory loss, we performed EEG spectral analyses over the entire break duration and explored how the power spectrum varied as a function of the group and memory performance (see ***Methods***). We first confirmed the expected between-groups difference in power spectral profiles (**Figure 3A**) and alpha/theta ratios (a common marker of sleepiness, **Figure 3B**), both of which showed a gradual increase in sleep depth between Wake, N1, and N2 subjects (alpha power and alpha/theta ratio differed between groups; mean alpha/theta ratio for Wake = 1.42, N1 = 1.21, N2 = 1.03; F(2,49) = 21.12, p<0.001, One-way ANOVA). Second and more importantly, a model comparison (see **Methods**) showed that the model best explaining forgetting included both the subject group and the alpha/theta ratio (comparison with the model including only the group information: F(1) = 4.63; p = 0.037). Considering this winning model, post-hoc comparisons indicated that both the alpha/theta ratio and the subject group were predictive of the forgetting rate (F(1) = 5.54, p = 0.038; F(2) = 5.52, p = 0.0069). To isolate the effect of the alpha/theta ratio, we normalized (z-scored) the alpha/theta ratios and forgetting rates within each group and quantified the correlation between these variables. We found a significant negative correlation between the alpha/theta ratio and the forgetting rate in all sleepers (including both the N1 and N2 groups; **Figure 3C**; r = −0.39; p = 0.019, Pearson correlation), indicating that the more participants’ brain activity slowed down during the break (lower alpha/theta ratio), the more they forgot the pictures’ location. Of note, this correlation did not reach significance (only a trend) when Wake subjects were added (**Supplementary Figure S2**; r = −0.25; p = 0.071, Pearson correlation). Overall, these EEG results are consistent with our behavioral results, since a low alpha/theta ratio (a good indicator of the N1 stage) was linked with a high forgetting rate. But they also extend the behavioral findings in two significant ways. First, they demonstrate that variations in the alpha/theta levels influence the extent of memory loss even within a single substage such as N1. Second, they provide a neurophysiological index for memory forgetting independent of the sleep stages.

**Figure 3:**
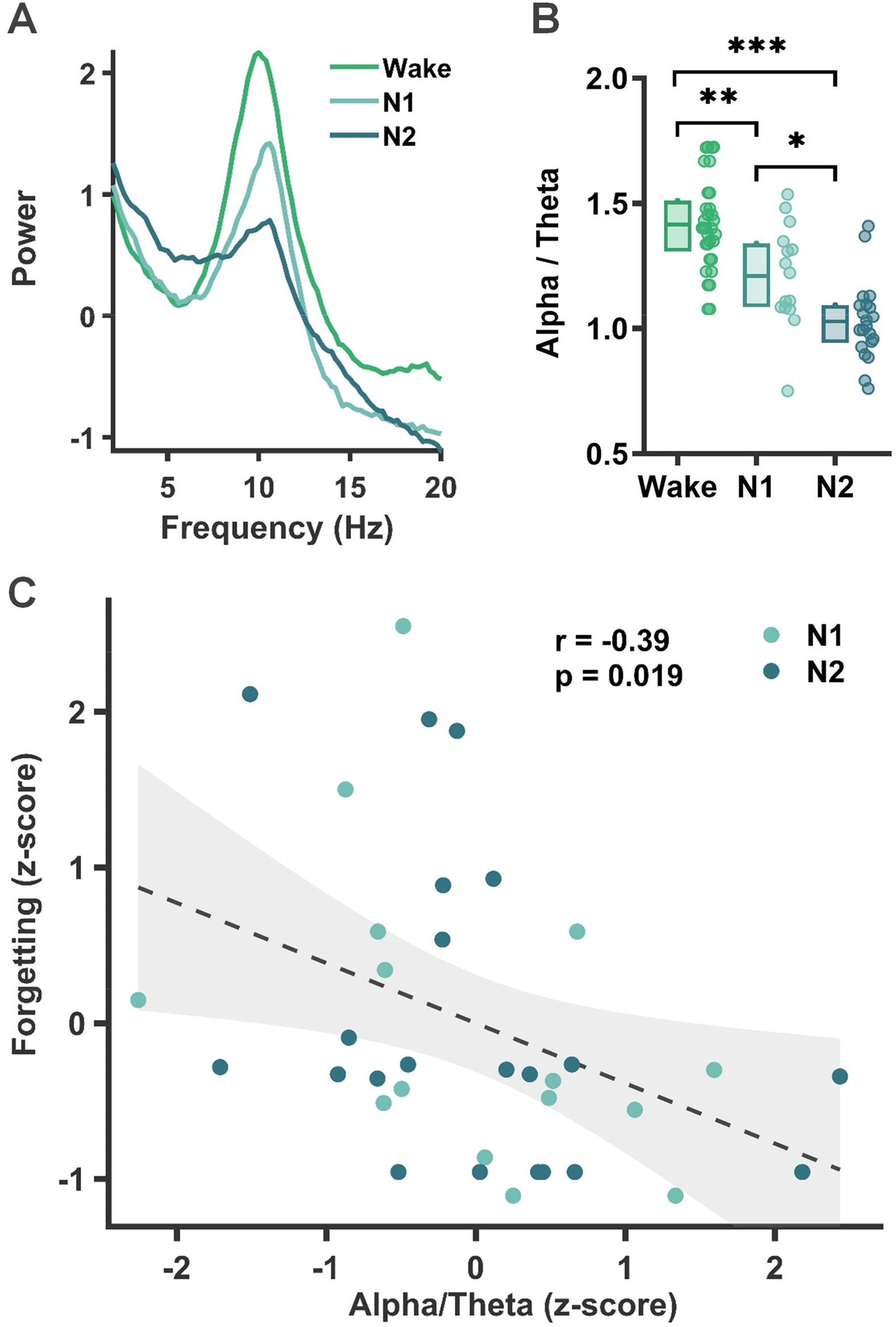
Neurophysiological substrate of memory loss. **(A)** Average power spectrum and **(B)** alpha/theta ratio over the occipital electrode during the Break for each group (Wake, N1, and N2). *** p<0.001; **, p<0.01; *, p<0.05 (One-way ANOVA for comparisons between all three groups and Tukey-Kramer post-hoc comparisons with correction for multiple comparisons). **(C)** Pearson correlation between the forgetting rate (z-score) and the alpha/theta ratio (z-score) for all the subjects who slept (N1 and N2 groups). Both the raw individual data (circles) and a glm fit with a 95% confidence interval (line+shaded area) are plotted. Rho and p-values are displayed in the figure.

## DISCUSSION

Countless studies on the relationship between sleep and memory have flourished in recent decades (Born et al., 2006; Diekelmann & Born, 2010; Rasch & Born, 2013). However, most of these studies focused on the role of the NREM sleep stages N2 and N3, and, to a lesser extent, of REM sleep, but left aside the specific impact of the first stage of NREM sleep (N1) on memory processing. Instead, most studies mix N1 sleep with the following NREM sleep stages or with wakefulness. We overcame this issue by devising an experimental paradigm allowing us to isolate the specific role of N1 sleep in the fate of recently encoded memories. We found that a resting period including only N1 sleep was associated with a lower memory recall (an average loss of 10% of previously encoded items) compared to a resting period including both N1 and N2. Of note, this memory loss was driven by the items that subjects struggled the most to encode during learning. It should be noted that this memory loss was driven by the items that the subjects struggled the most to encode during learning.

We extended this finding with EEG spectral analyses, which showed a negative correlation between the ratio of alpha/theta power (a marker of alertness) and the forgetting rate. This means that, for both N1 and N2 subjects, a deeper sleep (lower alpha/theta ratio) was associated with a higher forgetting rate. This result could seem at odds with the increase in forgetting observed in N1 compared to N2. However, it could be explained by the involvement of neural mechanisms specific to N2 (e.g., sleep spindles, K-complexes, and slow-waves, absent in N1 sleep) that are not captured by the alpha/theta ratio and may promote memory consolidation. Moreover, the alpha/theta ratio could represent a common physiological index of memory loss that is complementary to sleep stages and easier to extract.

Our finding that N1 sleep is associated with forgetting is consistent with the findings of two previous studies (Wyatt et al., 1994, 1997), which reported a decrease in recall of stimuli presented a few minutes prior to sleep onset. Yet, their paradigm differed from the present one in that their subjects were presented with stimuli while falling asleep, whereas in our study, participants had already encoded memories before reaching N1 sleep. Therefore, one may question whether the observed memory deficit in those 1990s studies was related to an active role of the sleep-onset period in forgetting or to reduced cognitive abilities (e.g., decreased encoding) associated with the alertness decline accompanying sleep onset (Ogilvie, 2001). The Wyatt results may thus reflect a deficiency in encoding, whereas our study shows the forgetting of correctly encoded items.

Of note, the between-group differences in memory performance observed in this study are unlikely related to sleep inertia at awakening from N1 or N2 sleep, as all groups had comparable alertness levels at the beginning of the *Post* phase (see **Table 1**). Plus, we did not find a similar memory loss in N2 participants (who are more likely to experience sleep inertia upon awakening from N2 sleep than the subjects awakening from N1 sleep). Overall, our findings suggest that N1 sleep specifically yields to the forgetting of recently encoded memories, particularly the ones that were encoded with the greatest difficulty.

At first glance, our results appear to contradict those of another study (Lahl et al., 2008), which reported enhanced retention of word lists following a 6-min ultra-short nap. In that study, however, the short nap was long enough to include N2 episodes, and the authors lumped together N1 and N2 sleep. This distinction with our paradigm is crucial since N1 and N2 might have opposing effects on memory. We found that subjects who slept only in N1 had a larger memory impairment than subjects who continued into deeper N2 sleep. However, in contrast with an extensive literature indicating a beneficial role of N2 sleep in memory consolidation, we did not observe any difference in memory performance between the Wake and N2 groups. One could argue that the periods of N2 sleep in our study were ‘lighter’ than those observed during longer naps. Indeed, although our participants were allowed to sleep for a maximal duration of 30 minutes, they spent only a few minutes in N2 (and not necessarily continuously), as they were repeatedly awakened during their naps. Thus, it is possible that a more consolidated and uninterrupted period of sleep, including a higher density of spindles and slow oscillations, is required to observe memory benefits (Cairney et al., 2018; Marshall et al., 2006; Mednick et al., 2013; Ngo et al., 2013). It is worth noting at this point that N1 data is sometimes lumped together with wake data in previous studies investigating memory consolidation. In that case, the authors would obtain comparable results to ours, namely a higher forgetting rate in their ‘wake’ group (possibly driven by N1) than in the N2 group. Therefore, this complementary hypothesis may also partially explain why we did not observe the common N2 sleep benefit on memory.

What role does N1 sleep play in sleep-related memory processing? Based on our findings, we see two possibilities. First, in accordance with the synaptic downscaling hypothesis (Tononi & Cirelli, 2006, 2014), N1 sleep may be involved in the active suppression of items with a weaker synaptic weight (the Lost object in our study). Such erasure of information could be necessary to make room in the synaptic network for subsequent memory consolidation in N2 sleep. Besides, forgetting is viewed as an essential function of sleep for efficient learning, complementary to memory consolidation (Feld & Born, 2017; Hardt et al., 2013; Poe, 2017). A few studies indicate that sleep actively contributes to forgetting by preferentially consolidating some information and pruning out others (Payne et al., 2008). Additionally, Feld et al. (2016) showed a sleep-dependent forgetting effect when the amount of encoded information was large. The authors hypothesized that large-scale data encoding results in overlapping hippocampal representations (Feld et al., 2016). During the following sleep, the gist (overlap) would then be preferentially consolidated, and the memory traces would be pruned out following global synaptic renormalization. This process could occur during N1 sleep and account for the observed forgetting of a few items.

An alternative hypothesis would be that N1 sleep instead initiates memory reprocessing by tagging the memory traces most in need of being reinforced (the weaker ones) by subsequent sleep stages. In that case, by preventing deeper sleep from supplanting N1, we would have aborted this process, resulting in the forgetting of these peculiar items (potentially by destabilizing those memory traces; Bonin & Koninck, 2015). Further studies would be needed to disentangle between these hypotheses, for example, by using multiple short naps of different natures with memory tests performed between them (e.g., testing whether a second nap including N2 sleep rescues memories that had been forgotten after a first nap including only N1 sleep). Whichever hypothesis turns out to be correct, the present findings indicate that some kind of memory processing occurs at sleep onset and highlight two different patterns between two seemingly proximal sleep stages, N1 and N2. Interestingly, these findings parallel those we recently observed about creativity (Lacaux et al., 2021). Indeed, we discovered that N1 sleep was associated with a boost in insight (i.e., the sudden discovery of a hidden regularity in the task), but this benefit vanished if subjects reached N2 sleep. This analogy suggests that 1) memory and creativity may be intertwined, and 2) it is important to consider N1 and N2 sleep stages separately in future studies.

Unfortunately, we did not have enough subjects who reported unambiguous task-related dreams to evaluate their impact on subsequent performance. Whether task-related dreaming directly impacts memory performance (as implied by some studies; Wamsley, Tucker, et al., 2010; Wamsley & Stickgold, 2019) or is merely an epiphenomenal reflection of ongoing memory reprocessing remains an open question.

In conclusion, by using a novel design that separates N1 from N2 sleep, we discovered that memory processing also takes place during N1 sleep, a stage that appears to promote memory forgetting. Interestingly, this function was originally hypothesized to be associated with another sleep stage, REM sleep (Crick & Mitchison, 1983), a theory that has recently received empirical support (Izawa et al., 2019; Li et al., 2017). These latest studies provide a putative mechanism underlying forgetting via the pruning of certain synapses, a mechanism that could thus occur as early as during N1 sleep. We hope that our work will launch further investigations to corroborate such results and better understand the mechanisms at work during N1 sleep and its role in determining the fate of our memories.

## Data availability

All the relevant data is available upon reasonable request. Inquiries should be directed to the corresponding author.

## Acknowledgements

This research was supported by the doctoral school ED3C and by the SFRMS (Société Française de Recherche et Médecine du Sommeil; grants to CL), as well as by Inserm (Institut National de la Santé et de la Recherche Médicale; research funds to DO). We thank C. Bastoul and A. Le Coz for helping acquire some data, S. Leu-Semenescu for sleep stage coding, and J. Frain for helping pilot the experimental task.

## Author contributions

CL and DO designed the study. CL collected the data. CL and TA analyzed the data. CL, TA, and DO wrote the paper. IA furnished the study infrastructure. DO and IA supervised the study.

## References

Berry, R. B., Brooks, R., Gamaldo, C. E., Harding, S. M., Marcus, C., & Vaughn, B. V. (2012). The AASM manual for the scoring of sleep and associated events. Rules, Terminology and Technical Specifications, Darien, Illinois, American Academy of Sleep Medicine, 176, 2012.

Bonin, R. P., & Koninck, Y. D. (2015). Reconsolidation and the regulation of plasticity: Moving beyond memory. Trends in Neurosciences, 38(6), 336–344.

Born, J., Rasch, B., & Gais, S. (2006). Sleep to remember. The Neuroscientist: A Review Journal Bringing Neurobiology, Neurology and Psychiatry, 12(5), 410–424.

Brainard, D. H. (1997). The Psychophysics Toolbox. Spatial Vision, 10(4), 433–436.

Cairney, S. A., Guttesen, A. Á. V., El Marj, N., & Staresina, B. P. (2018). Memory Consolidation Is Linked to Spindle-Mediated Information Processing during Sleep. Current Biology: CB, 28(6), 948–954.e4.

Crick, F., & Mitchison, G. (1983). The function of dream sleep. Nature, 304(5922), 111–114.

De Koninck, J., Christ, G., Hébert, G., & Rinfret, N. (1990). Language learning efficiency, dreams and REM sleep. Psychiatric Journal of the University of Ottawa: Revue De Psychiatrie De l ‘Universite d’Ottawa, 15(2), 91–92.

Diekelmann, S., & Born, J. (2010). The memory function of sleep. Nature Reviews. Neuroscience, 11(2), 114–126.

Feld, G. B., & Born, J. (2017). Sculpting memory during sleep: Concurrent consolidation and forgetting. Current Opinion in Neurobiology, 44, 20–27.

Feld, G. B., Weis, P. P., & Born, J. (2016). The Limited Capacity of Sleep-Dependent Memory Consolidation. Frontiers in Psychology, 7, 1368.

Giuditta, A. (2014). Sleep memory processing: The sequential hypothesis. Frontiers in Systems Neuroscience, 8.

Grant, D. A., Honn, K. A., Layton, M. E., Riedy, S. M., & Van Dongen, H. P. A. (2017). 3-minute smartphone-based and tablet-based psychomotor vigilance tests for the assessment of reduced alertness due to sleep deprivation. Behavior Research Methods, 49(3), 1020–1029.

Hardt, O., Nader, K., & Nadel, L. (2013). Decay happens: The role of active forgetting in memory. Trends in Cognitive Sciences, 17(3), 111–120.

Hertig-Godeschalk, A., Skorucak, J., Malafeev, A., Achermann, P., Mathis, J., & Schreier, D. R. (2020). Microsleep episodes in the borderland between wakefulness and sleep. Sleep, 43(1).

Hori, T., Hayashi, M., & Morikawa, T. (1994). Topographical EEG changes and the hypnagogic experience. In Sleep onset: Normal and abnormal processes (pp. 237–253). American Psychological Association.

Izawa, S., Chowdhury, S., Miyazaki, T., Mukai, Y., Ono, D., Inoue, R., Ohmura, Y., Mizoguchi, H., Kimura, K., Yoshioka, M., Terao, A., Kilduff, T. S., & Yamanaka, A. (2019). REM sleep–active MCH neurons are involved in forgetting hippocampus-dependent memories. Science, 365(6459), 1308–1313.

JASP Team. (2019). JASP (Version) [Computer software]. https://jasp-stats.org/

Kleiner, M., Brainard, D. H., Pelli, D., Ingling, A., Murray, R., & Broussard, C. (2007). What’s new in Psychtoolbox-3. Perception, 36, 1–16.

Lacaux, C., Andrillon, T., Bastoul, C., Idir, Y., Fonteix-Galet, A., Arnulf, I., & Oudiette, D. (2021). Sleep onset is a creative sweet spot. Science Advances.

Lacroix, M. M., Lavilléon, G. de, Lefort, J., Kanbi, K. E., Bagur, S., Laventure, S., Dauvilliers, Y., Peyron, C., & Benchenane, K. (2018, December 7). Improved sleep scoring in mice reveals human-like stages. 489005.

Lahl, O., Wispel, C., Willigens, B., & Pietrowsky, R. (2008). An ultra short episode of sleep is sufficient to promote declarative memory performance. Journal of Sleep Research, 17(1), 3–10.

Li, W., Ma, L., Yang, G., & Gan, W.-B. (2017). REM sleep selectively prunes and maintains new synapses in development and learning. Nature Neuroscience, 20(3), 427–437.

Maingret, N., Girardeau, G., Todorova, R., Goutierre, M., & Zugaro, M. (2016). Hippocampo-cortical coupling mediates memory consolidation during sleep. Nature Neuroscience, 19(7), 959–964.

Marshall, L., Helgadóttir, H., Mölle, M., & Born, J. (2006). Boosting slow oscillations during sleep potentiates memory. Nature, 444(7119), 610–613.

Mednick, S. C., McDevitt, E. A., Walsh, J. K., Wamsley, E., Paulus, M., Kanady, J. C., & Drummond, S. P. A. (2013). The Critical Role of Sleep Spindles in Hippocampal-Dependent Memory: A Pharmacology Study. The Journal of Neuroscience, 33(10), 4494–4504.

Morel, P. (2018). Gramm: Grammar of graphics plotting in Matlab. J. Open Source Softw.

Ngo, H.-V. V., Martinetz, T., Born, J., & Mölle, M. (2013). Auditory Closed-Loop Stimulation of the Sleep Slow Oscillation Enhances Memory. Neuron, 78(3), 545–553.

Ogilvie, R. D. (2001). The process of falling asleep. Sleep Medicine Reviews, 5(3), 247–270.

Ólafsdóttir, H. F., Bush, D., & Barry, C. (2018). The Role of Hippocampal Replay in Memory and Planning. Current Biology, 28(1), R37–R50.

Oudiette, D., Antony, J. W., Creery, J. D., & Paller, K. A. (2013). The Role of Memory Reactivation during Wakefulness and Sleep in Determining Which Memories Endure. Journal of Neuroscience, 33(15), 6672–6678.

Paller, K. A., Creery, J. D., & Schechtman, E. (2021). Memory and Sleep: How Sleep Cognition Can Change the Waking Mind for the Better. Annual Review of Psychology, 72(1), 123–150.

Payne, J. D., Stickgold, R., Swanberg, K., & Kensinger, E. A. (2008). Sleep preferentially enhances memory for emotional components of scenes. Psychological Science, 19(8), 781–788.

Picchioni, D., Fukunaga, M., Carr, W. S., Braun, A. R., Balkin, T. J., Duyn, J. H., & Horovitz, S. G. (2008). FMRI differences between early and late stage-1 sleep. Neuroscience Letters, 441(1), 81–85.

Plailly, J., Villalba, M., Vallat, R., Nicolas, A., & Ruby, P. (2019). Incorporation of fragmented visuo-olfactory episodic memory into dreams and its association with memory performance. Scientific Reports, 9(1), 15687.

Poe, G. R. (2017). Sleep Is for Forgetting. Journal of Neuroscience, 37(3), 464–473.

Rasch, B., & Born, J. (2013). About Sleep’s Role in Memory. Physiological Reviews, 93(2), 681–766.

Rudoy, J. D., Voss, J. L., Westerberg, C. E., & Paller, K. A. (2009). Strengthening Individual Memories by Reactivating Them During Sleep. Science, 326(5956), 1079–1079.

Saletin, J. M., Goldstein, A. N., & Walker, M. P. (2011). The role of sleep in directed forgetting and remembering of human memories. Cerebral Cortex (New York, N.Y.: 1991), 21(11), 2534–2541.

Schneider, S. L. (2013). The International Standard Classification of Education 2011. In Class and Stratification Analysis (Vol. 30, pp. 365–379). Emerald Group Publishing Limited.

Shohamy, D., & Wagner, A. D. (2008). Integrating memories in the human brain: Hippocampal-midbrain encoding of overlapping events. Neuron, 60(2), 378–389.

Stenstrom, P., Fox, K., Solomonova, E., & Nielsen, T. (2012). Mentation during sleep onset theta bursts in a trained participant: A role for NREM stage 1 sleep in memory processing? International Journal of Dream Research, 5(1), 37–46.

Stickgold, R. (2009). How Do I Remember? Let Me Count the Ways. Sleep Medicine Reviews, 13(5), 305–308.

Stickgold, R., Malia, A., Maguire, D., Roddenberry, D., & O’Connor, M. (2000). Replaying the game: Hypnagogic images in normals and amnesics. Science (New York, N.Y.), 290(5490), 350–353.

Stickgold, R., & Walker, M. P. (2013). Sleep-dependent memory triage: Evolving generalization through selective processing. Nature Neuroscience, 16(2), 139–145.

Tononi, G., & Cirelli, C. (2006). Sleep function and synaptic homeostasis. Sleep Medicine Reviews, 10(1), 49–62.

Tononi, G., & Cirelli, C. (2014). Sleep and the price of plasticity: From synaptic and cellular homeostasis to memory consolidation and integration. Neuron, 81(1), 12–34.

Wamsley, E. J., Perry, K., Djonlagic, I., Reaven, L. B., & Stickgold, R. (2010). Cognitive replay of visuomotor learning at sleep onset: Temporal dynamics and relationship to task performance. Sleep, 33(1), 59–68.

Wamsley, E. J., & Stickgold, R. (2019). Dreaming of a learning task is associated with enhanced memory consolidation: Replication in an overnight sleep study. Journal of Sleep Research, 28(1), e12749.

Wamsley, E. J., Tucker, M., Payne, J. D., Benavides, J. A., & Stickgold, R. (2010). Dreaming of a learning task is associated with enhanced sleep-dependent memory consolidation. Current Biology: CB, 20(9), 850–855.

Wyatt, J. K., Bootzin, R. R., Allen, J. J. B., & Anthony, J. L. (1997). Mesograde Amnesia During the Sleep Onset Transition: Replication and Electrophysiological Correlates. Sleep, 20(7), 512–522.

Wyatt, J. K., Bootzin, R. R., Anthony, J., & Bazant, S. (1994). Sleep onset is associated with retrograde and anterograde amnesia. Sleep, 17(6), 502–511.

