## Supplementary Figures for "Memory loss at sleep onset"

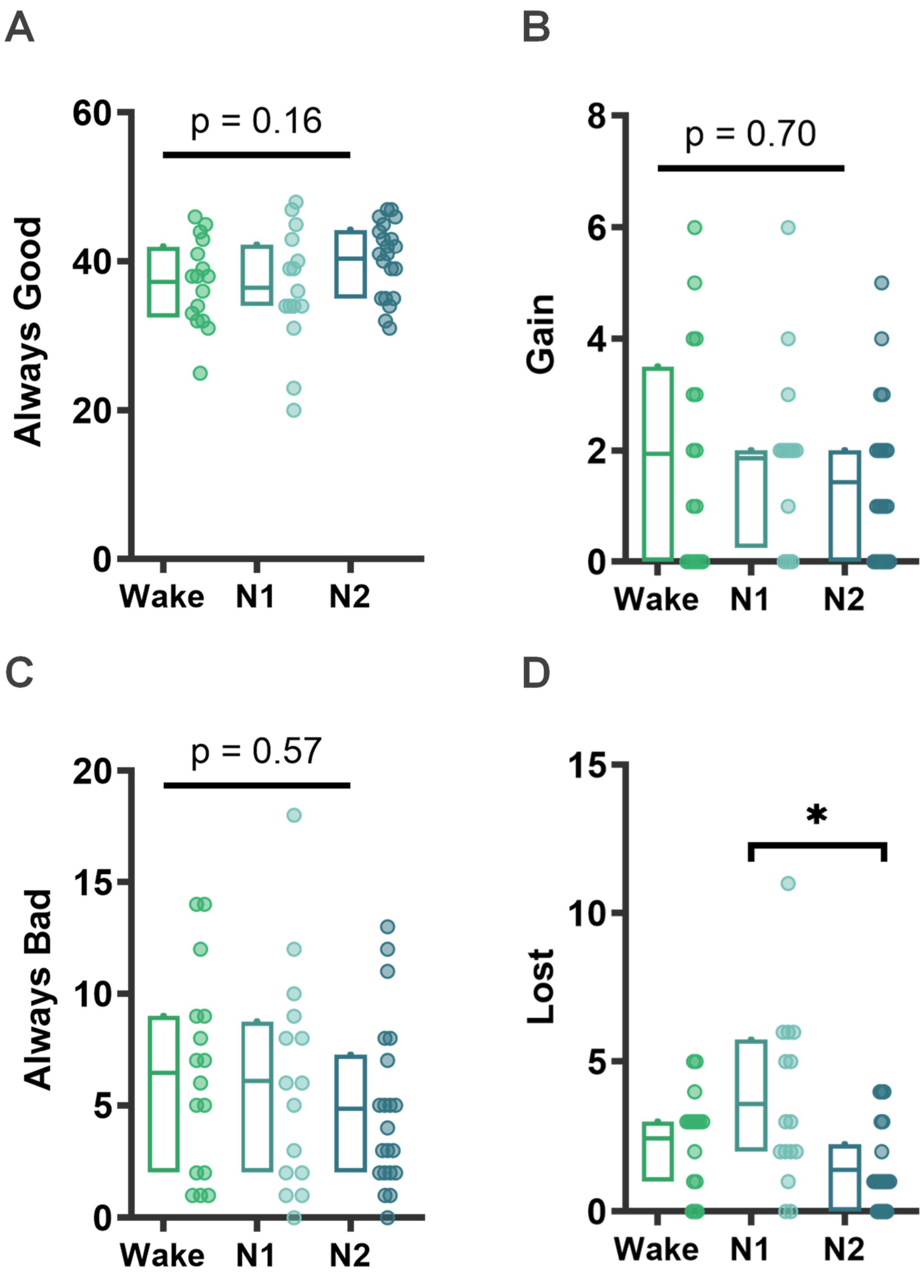


***Figure S1 - Repartition of the objects.*** *Number of objects in each of the 4 defined categories for all groups: 1) Always Good: objects correctly located in both the Pre and Post phases, 2) Gain: objects falsely located in the Pre phase but correctly located in the Post phase, 3) Always Bad: false in both the Pre and Post phases, and 4) Lost: correctly located in Pre but falsely located in Post. *p<0.05, n.s., for non-significant differences between groups (Kruskal-Wallis).*

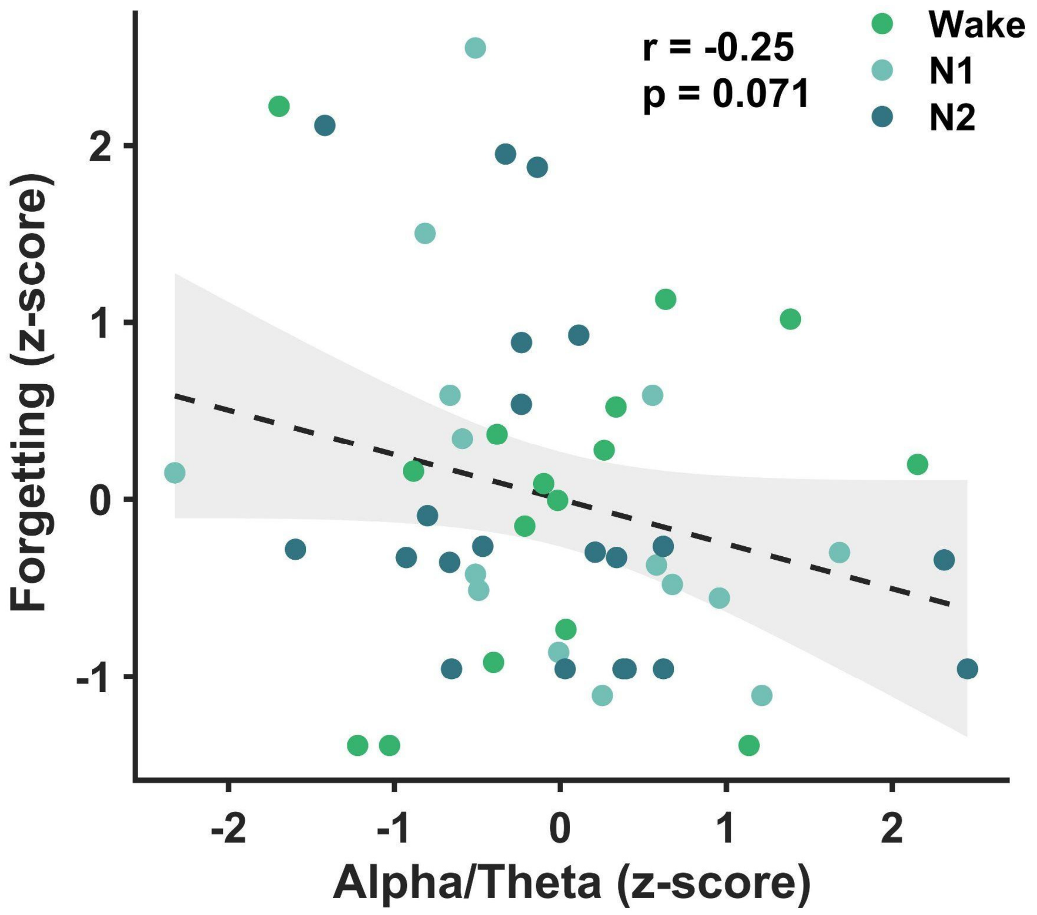


***Figure S2 -  Pearson correlation between forgetting rate (z-score) and alpha/theta ratio (z-score)*** *for all subjects (Wake, N1 and N2 groups). Both the raw individual data (circles) and a glm fit with a 95% confidence interval (line+shaded area) are plotted. Rho and p-values are displayed in the figure.*
